## Supplementary figures and images for "A cerebellar internal model calibrates a feedback controller involved in sensorimotor control"

### Supplementary Figure 1

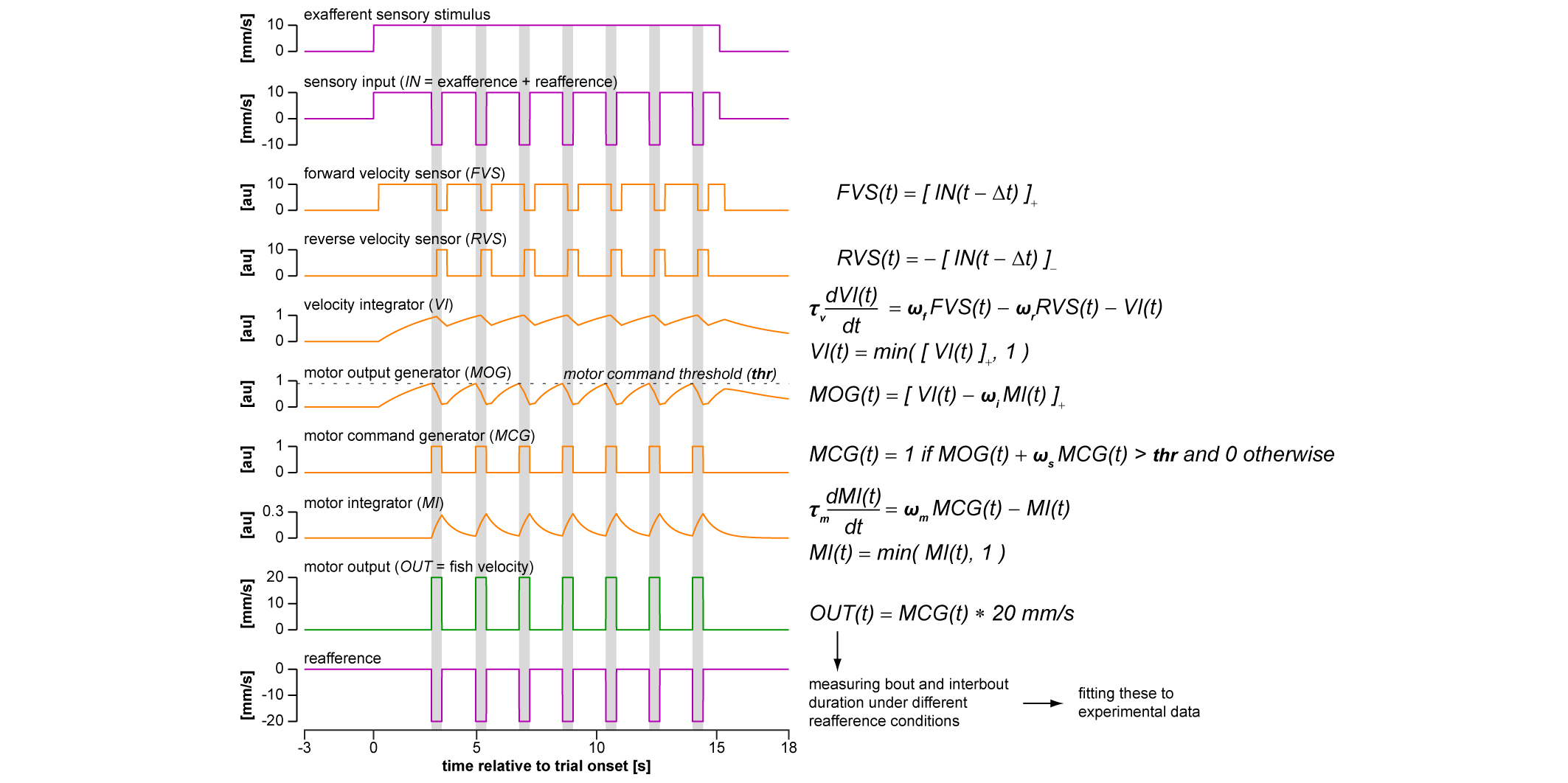

### Supplementary Figure 2

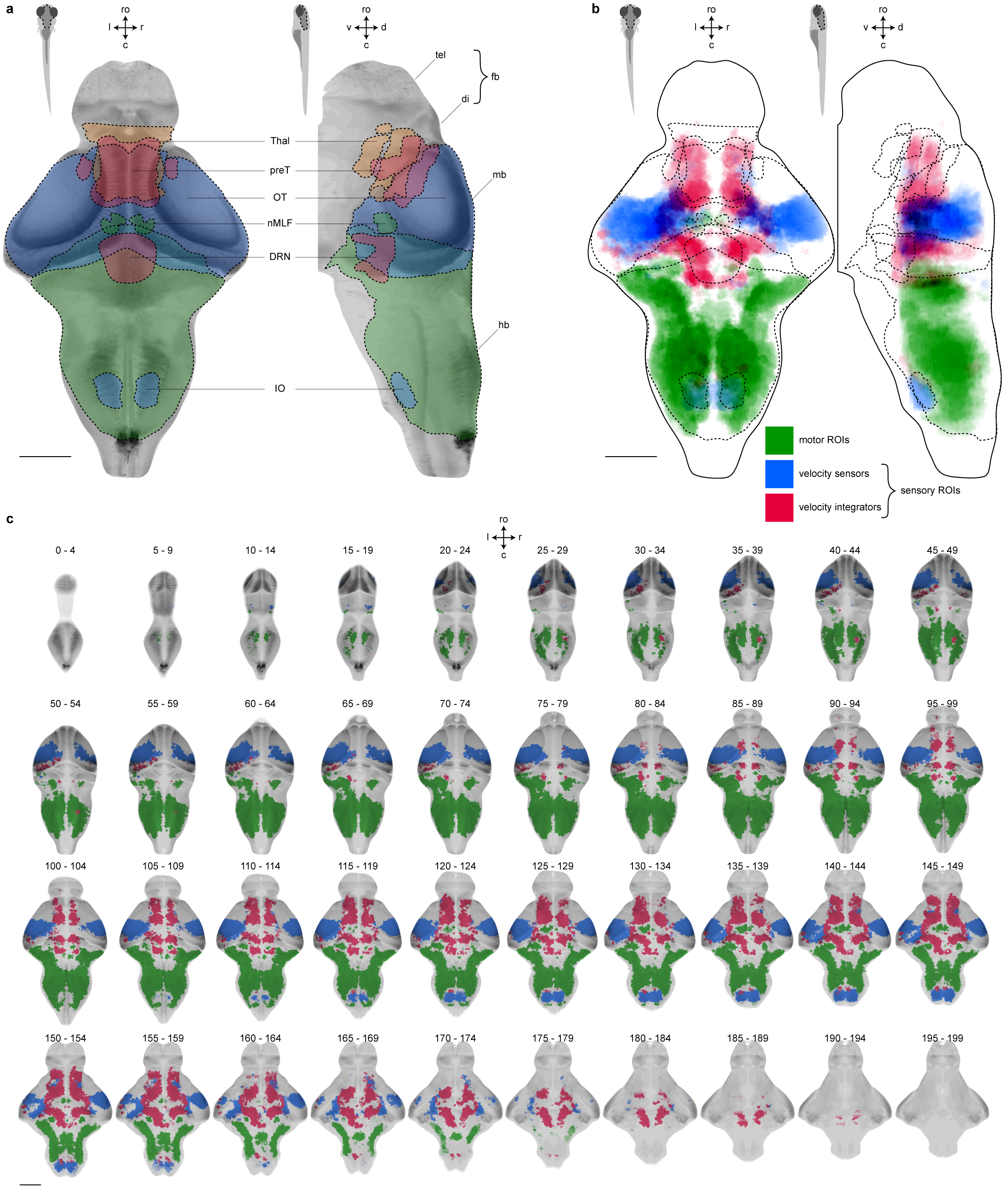

### Supplementary Figure 3

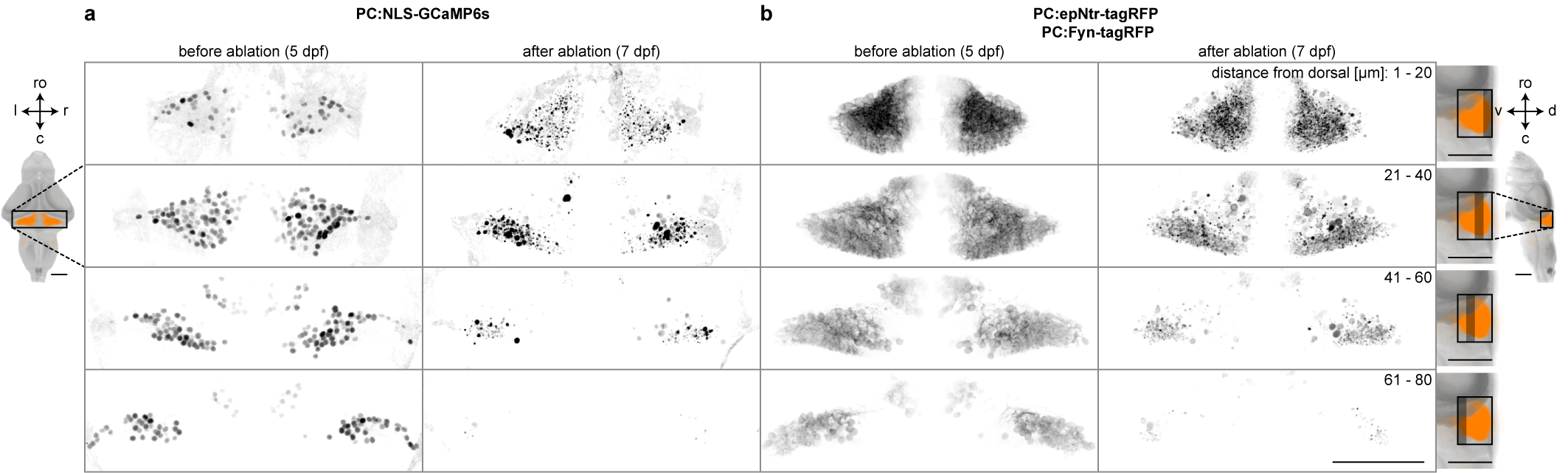

### Supplementary Figure 4

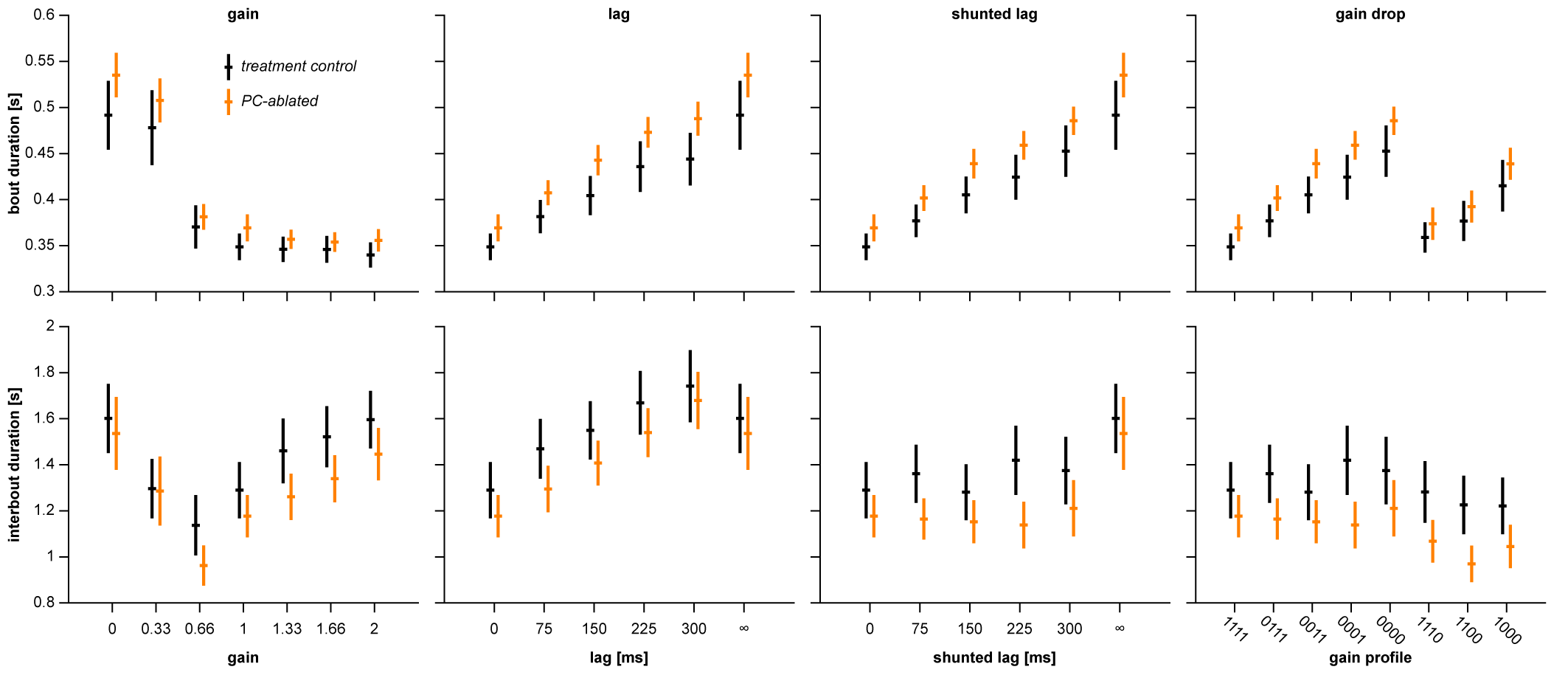

### Supplementary Figure 5

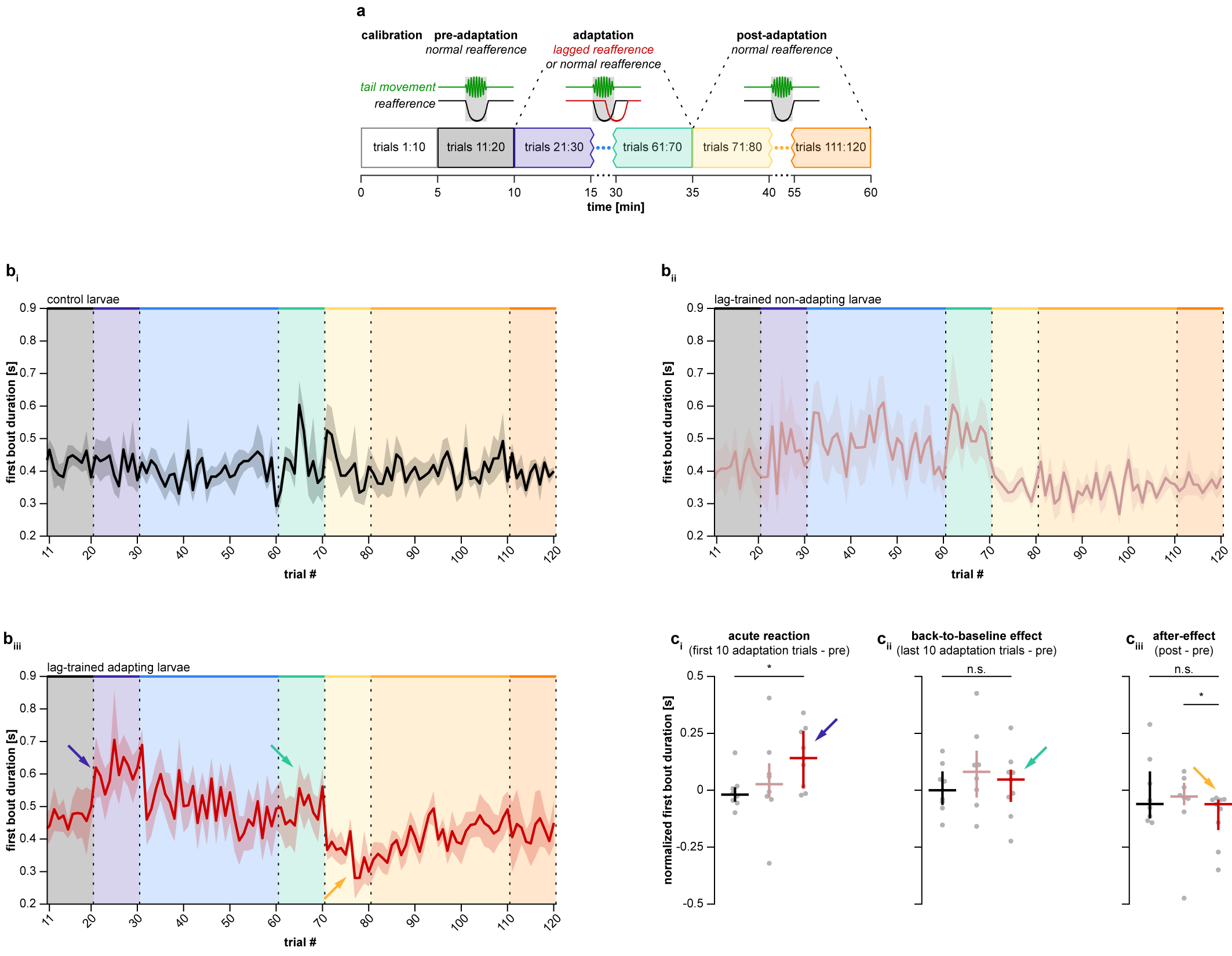

### Supplementary Figure 6

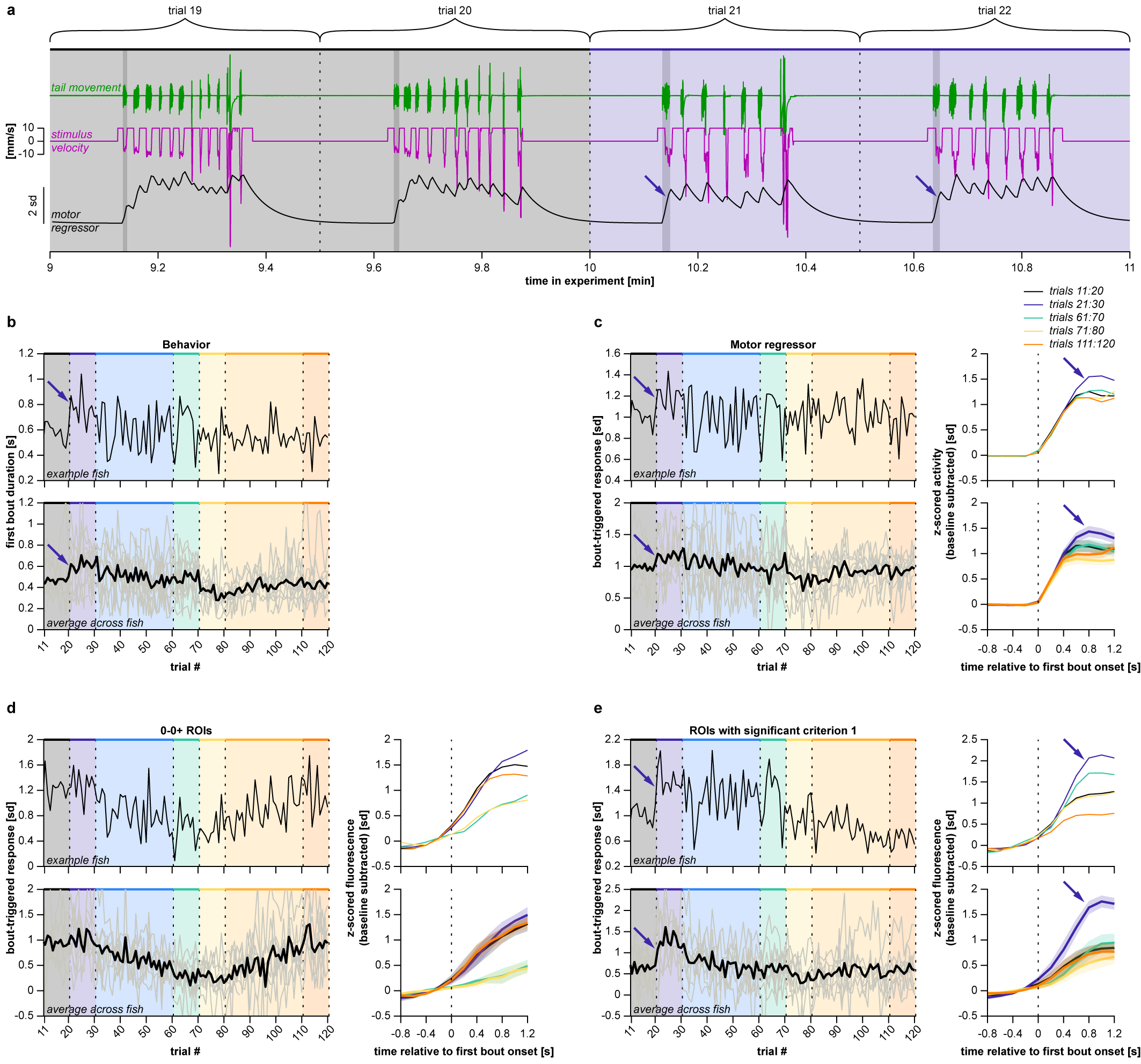

### Supplementary Figure 7

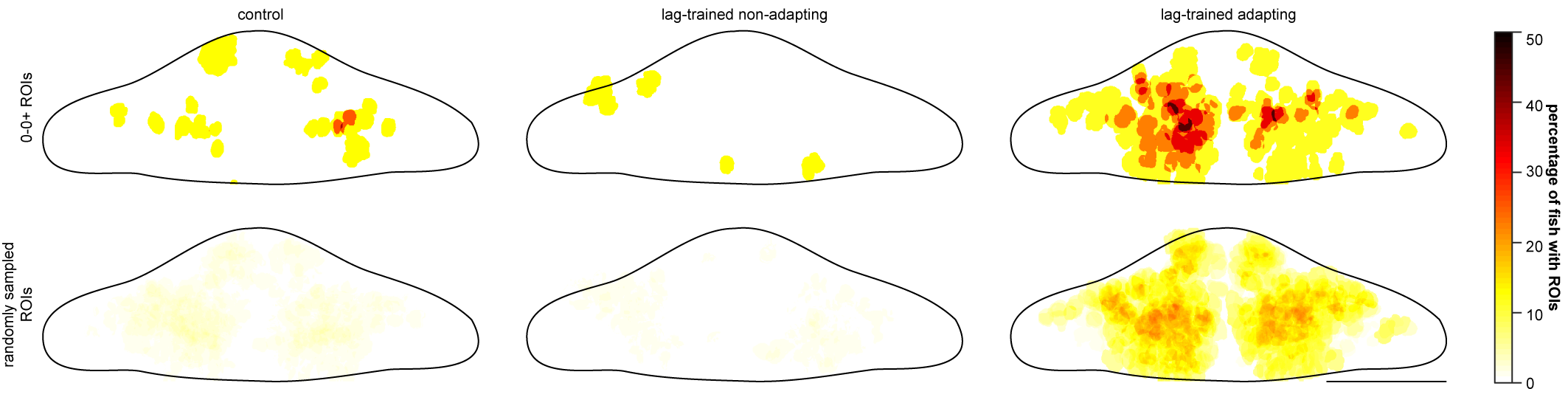
